# South American freshwater fish diversity shaped by Andean uplift since the Late Cretaceous

**DOI:** 10.1101/2021.05.14.444133

**Authors:** Lydian M. Boschman, Fernanda A.S. Cassemiro, Luca Carraro, Jorad de Vries, Florian Altermatt, Oskar Hagen, Carina Hoorn, Loïc Pellissier

## Abstract

South America is home to the highest freshwater fish biodiversity on Earth^1,2^. The hotspot of species richness is located in the western Amazon Basin, and richness decreases downstream along the Amazon River towards the mouth at the Atlantic coast (Fig. 1b, c)^3,4^, which contradicts the positive relationship between stream size and biodiversity that is commonly observed in river systems across the world^5,6^. We investigate the role of river rerouting events caused by Andean mountain building and repeated episodes of flooding in western Amazonia in shaping the modern-day richness pattern of freshwater fishes in South America. To this end, we combine a reconstruction of river networks following Andean surface uplift since 80 million years ago with a mechanistic biological model simulating dispersal, allopatric speciation and extinction over the dynamic landscape of rivers and lakes. We show that the numerous small river rerouting events in western Amazonia resulting from mountain building produced highly dynamic riverine habitats that caused high diversification rates, shaping the exceptional present-day richness of this region. The history of marine incursions and lakes, including the Miocene Pebas megawetland system in western Amazonia, played a secondary role. This study is a major step towards the understanding of the processes involved in the interactions between the solid Earth, landscapes, and life of extraordinary biodiverse South America.

---

The commonly observed gradient of increasing biodiversity downstream a river is caused by the inherent properties of dendritic river networks, which drive connectivity and shape both present-day biotic and abiotic conditions in riverine landscapes^7-9^. But despite often being regarded as such, dendritic connectivity is not a static property. Deep-time paleogeographic and sealevel changes can drastically alter the structure and connectivity of river systems through river rerouting events, river captures, or drainage reversals. As a result, the highly dynamic deep-time paleogeographic history of South America, including the gradual uplift of the Andes since ∼100 million years ago (Ma) and the repeated flooding of large areas of the Amazon Basin, is expected to have left an imprint on the richness pattern observed today^3^. However, net effects of river reorganizations on biodiversity are complex: river captures merge previously isolated habitats and populations, thereby facilitating dispersal and geographical range expansion, which increases local diversity initially, but decreases rates of speciation and extinction. At the same time, river captures separate previously connected habitats, leading to an increase in genetic isolation and an increase in rates of speciation and extinction^10,11^. Additionally, the formation of lakes leads to the addition of habitat and the merging of populations, while the disappearance of lakes entails the opposite effects.

Phylogenetic and paleontological datasets indicate that the fish fauna of the Amazon region originated during the Late Cretaceous, and that diversity in Amazonia is the result of a prolonged history of net diversification in which speciation rates exceeded extinction rates^10,12,13^. There is no evidence for significant changes in speciation rates through time nor evidence of major extinction events^13-15^. Considering the complexity of the effects of river reorganizations and the stability of speciation rates, the diversification history of South American freshwater fish fauna cannot be explained through classic correlative approaches aimed at linking speciation or extinction events to paleogeographical events. Instead, it requires an approach in which the ambiguous effects of river reorganizations can be assessed in a process-based way, and in which paleogeographic changes are not considered to be ‘instantaneous’ (in the context of geological time, within a million year), but continuous. In this study, we use a mechanistic modelling approach^16^, simulating dispersal, allopatric speciation and extinction over a dynamic landscape of rivers and lakes. We investigate the role of Andean mountain building and lake system dynamics since 80 Ma in shaping the modern-day richness pattern of freshwater fishes in South America, and in particular, the enigmatic richness gradient in the Amazon Basin.

## Paleogeographic history of South America

Andean mountain building initiated in the Late Cretaceous, at ∼100 Ma in Patagonia, ∼80 Ma in the central Andes of Bolivia and Peru, and ∼70 Ma in the ranges of Ecuador^18^. During the Late Cretaceous and Paleocene, the northwestern corner of the South American continent was mostly covered by shallow seas^19^. The bulk of the northern South American rivers flowed towards this northwestern corner, collecting water and sediments from the Brazilian and Guianan shields and the incipient central Andes before draining into the Caribbean Sea and Pacific Ocean^20^. During the Eocene and Oligocene, uplift migrated to regions further north^21^ and a continuous continent-scale mountain range was established by the beginning of the Miocene, albeit significantly smaller in width and lower in elevation compared to the modern orogen^22^. The topography in the northern Andes that was established during the Oligocene-Miocene blocked drainage towards the Pacific, and additionally, resulted in the formation of a deep foreland basin^23^. As a result, the Miocene western Amazon Basin was characterized by a system of mountain-parallel rivers and wetlands named the Pebas system, which drained towards the north into the Caribbean Sea^20,24,25^. In eastern Amazonia, the precursor of the modern Amazon river was already present, draining into the Atlantic Ocean^24-26^. Continued uplift in the northern Andes in the Late Miocene and Pliocene produced erosional material forming ‘megafans’ along the eastern slopes of the mountains^27^. This erosion material gradually filled the Miocene foreland basins of western Amazonia, leading to the disappearance of the wetlands, and around ∼10 Ma, to the establishment of the transcontinental west-to-east flowing Amazon River^24-26^.

## Drainage network reconstruction

We generated river networks by combining a river reconstruction algorithm with a recently developed reconstruction of Andean mountain building since 80 Ma of Boschman^22^, consisting of a series of 80 paleo-digital elevation models (paleoDEMs, one per each million-year time step), at a 0.1° spatial resolution. The river reconstruction algorithm establishes drainage directions for every cell based on the steepest possible descent between neighboring grid cells, producing a realistic network of rivers on a given topography. Additionally, we incorporated the configurations of the Miocene Pebas wetlands system and other marine incursions and lakes based on Hoorn and Wesselingh^20^. We computed drainage networks for three paleogeographic scenarios designed to test the relative roles of the two main aspects of riverine landscape evolution in South America: (1) mountain building and the consequent river rerouting events, and (2) lake system dynamics, which includes the formation and disappearance of lakes (suitable habitat), and marine incursions (non-suitable habitat). Marine incursions modify the location of the shoreline, thereby altering the locations of river outlets. Scenario A includes lakes system dynamics but excludes mountain building (i.e. the topography is identical to modern topography in all times steps), scenario B includes mountain building but excludes lake system dynamics, and scenario C includes both mountain building and lake system dynamics.

Results from the river reconstruction algorithm (Fig. 2) indicate that drainage networks changed substantially throughout the last 80 Ma. However, these changes occurred primarily in the western Amazon Basin, whereas river networks in southern and eastern South America changed substantially less. Changes in habitat resulting from lake dynamics are larger but less frequent compared to the much more frequent but smaller changes due to mountain building (Fig. 2). Taking into account both surface uplift in the Andes and wetlands in the Amazon Basin (scenario C), the model produces river morphologies through time that accurately match first order river reconstruction from sedimentological and paleontological data as summarized above.

**Fig. 1.**
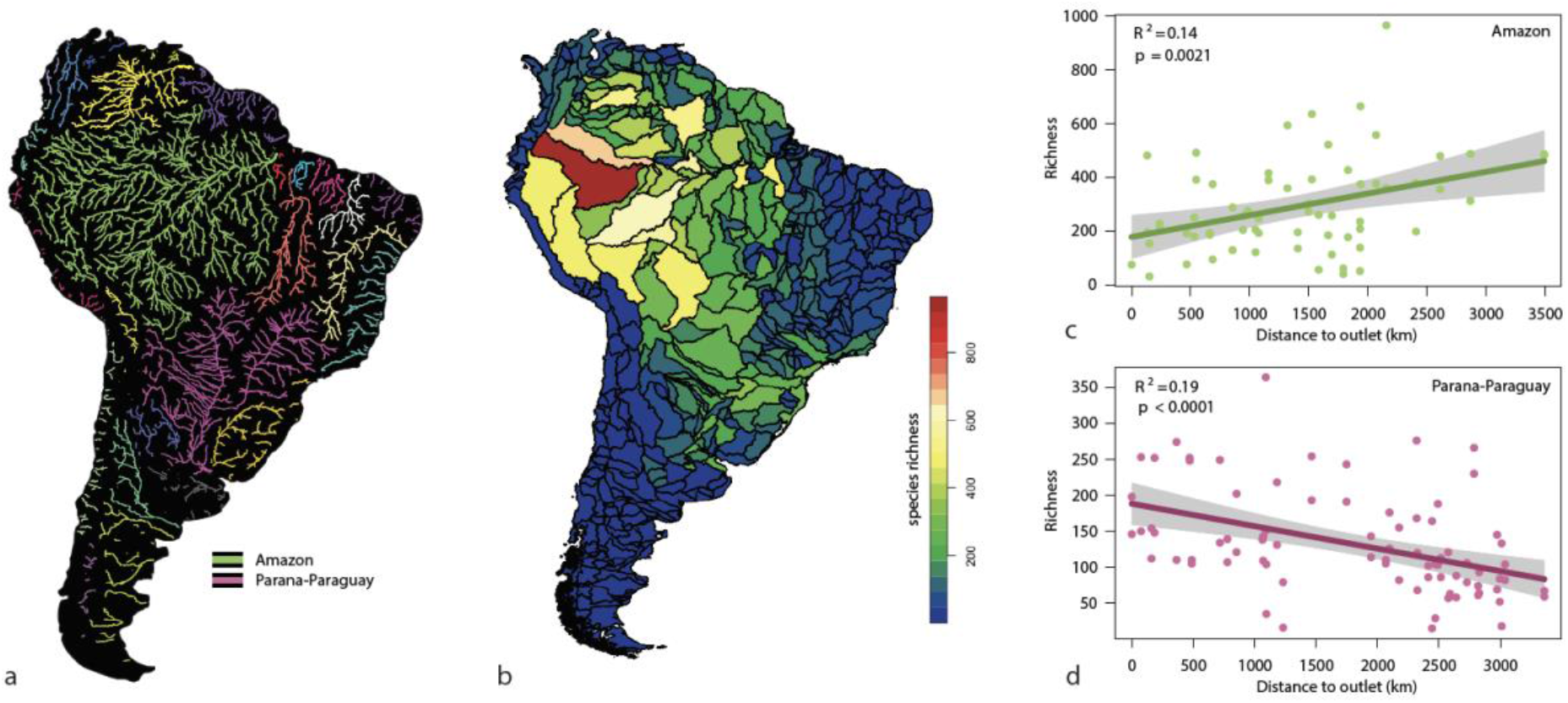
**a**, Major rivers and drainage basins of South America. HydroRIVERS data from the HydroSHEDS database^17^; rivers shown that drain a surface area of >4000 km^2^. **b**, Freshwater fish species richness per sub-basin (level 5 HydroBASINS^17^), reproduced from data from Cassemiro, et al. ^4^. **c and d**, Richness versus distance to outlet per sub-basins (as shown in panel b), illustrating the positive relationship (richness decreases towards the mouth of the river) for the Amazon (c), and, for comparison, a negative relationship for the Parana-Paraguay river basin (d).

**Fig. 2.**
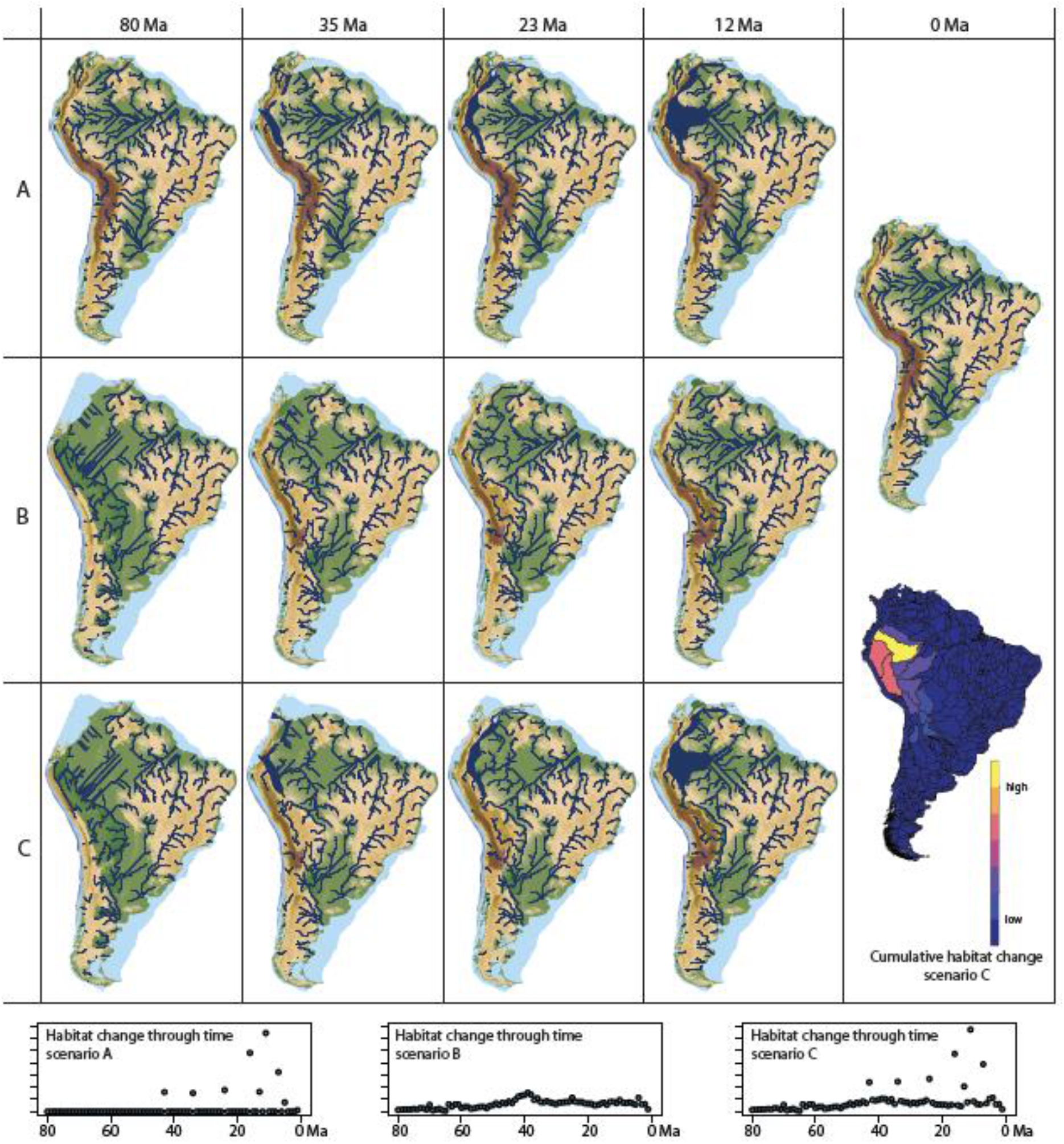
River networks for the three paleogeographic scenarios. A: Excluding mountain building, including lake system dynamics; B: including mountain building, excluding lake system dynamics; C: including both mountain building and lake system dynamics. In scenario A, the transcontinental west-to-east flowing Amazon river is present in all time steps, except for during the culmination of the Pebas wetlands system (15-11 Ma), when western Amazonia drained into the Pebas lake, and subsequently, into the Caribbean Sea. Scenarios B and C are identical for early time steps (80-43 Ma), before the presence of marine incursions and lakes, and very similar for 42-24 Ma, in which western Amazonia drained towards the north(west), and eastern Amazonia towards the east. Scenarios B and C start deviating significantly from 23 Ma onwards. In scenario B, the transcontinental west-to-east flowing Amazon river is established at 21 Ma. In scenario C, western Amazonia remains to drain towards the north(west) for 23-11 Ma, and only at 10 Ma, after the disappearance of the Pebas system, is the transcontinental Amazon river established. Habitat change represents the number of grid cells changing from suitable to unsuitable or vice versa from one time step to the next.

## Biodiversity dynamics

Using the mechanistic biodiversity model *gen3sis*^16^, we simulate biodiversity dynamics in a geographical framework through three mechanisms: dispersal, speciation, and extinction. The dispersal rate (distance per time step) determines how far species populations disperse through suitable habitat cells. Speciation occurs when two populations of the same species become disconnected through changes in habitat connectivity. This simplification of speciation (i.e. only through allopatry) is warranted, as assemblages of South American fish species are known to be polyphyletic, meaning that locally coexisting species are rarely each other’s close relatives and geographic ranges of sister species rarely overlap^28-35^. This implies that the origin of these species can primarily be attributed to allopatric speciation, i.e., the evolution of a new species as a result of isolation of two populations of a species, and that sympatric speciation, i.e., the evolution of a new species from a surviving ancestral species while both inhabit the same geographical range, played a negligible role in the diversification history of South American fishes^28,33,36,37^. Extinction occurs when all cells in the range of a species change from suitable (river or lake) to unsuitable habitat (land or marine).

Simulations of biodiversity dynamics on the river landscape generated by the river reconstruction algorithm (Fig. S1) consistently produced richness patterns with a hotspot in western Amazonia, irrespective of model parameters (Fig. S2-4) and paleogeographic input scenario (Fig. 3). We therefore conclude that the enigmatic richness gradient along the Amazon river can be explained by the paleogeographic history of South America. The simulation results indicate that the high species richness in western Amazonia can be attributed to high speciation rates caused by either the gradual uplift of the central and northern Andes or the history of flooding producing wetlands and lakes in western Amazonia, or both. Interestingly, speciation rates in scenario C are higher than the sum of speciation rates in scenario A and B (Fig. 3d-f), indicating that interactions between the two components of the paleogeographic history (mountain building and lake dynamics) intensify the diversification that can be explained by each component individually. However, the much higher number of speciation events resulting from the Andean uplift history (Fig. 3b) and the small difference in model output between scenarios B and C (Fig. 3b-c, e-f) indicate that the uplift of the Andes has been the primary driver of diversification of freshwater fishes in South America, while the history of flooding played a secondary role.

**Fig. 3.**
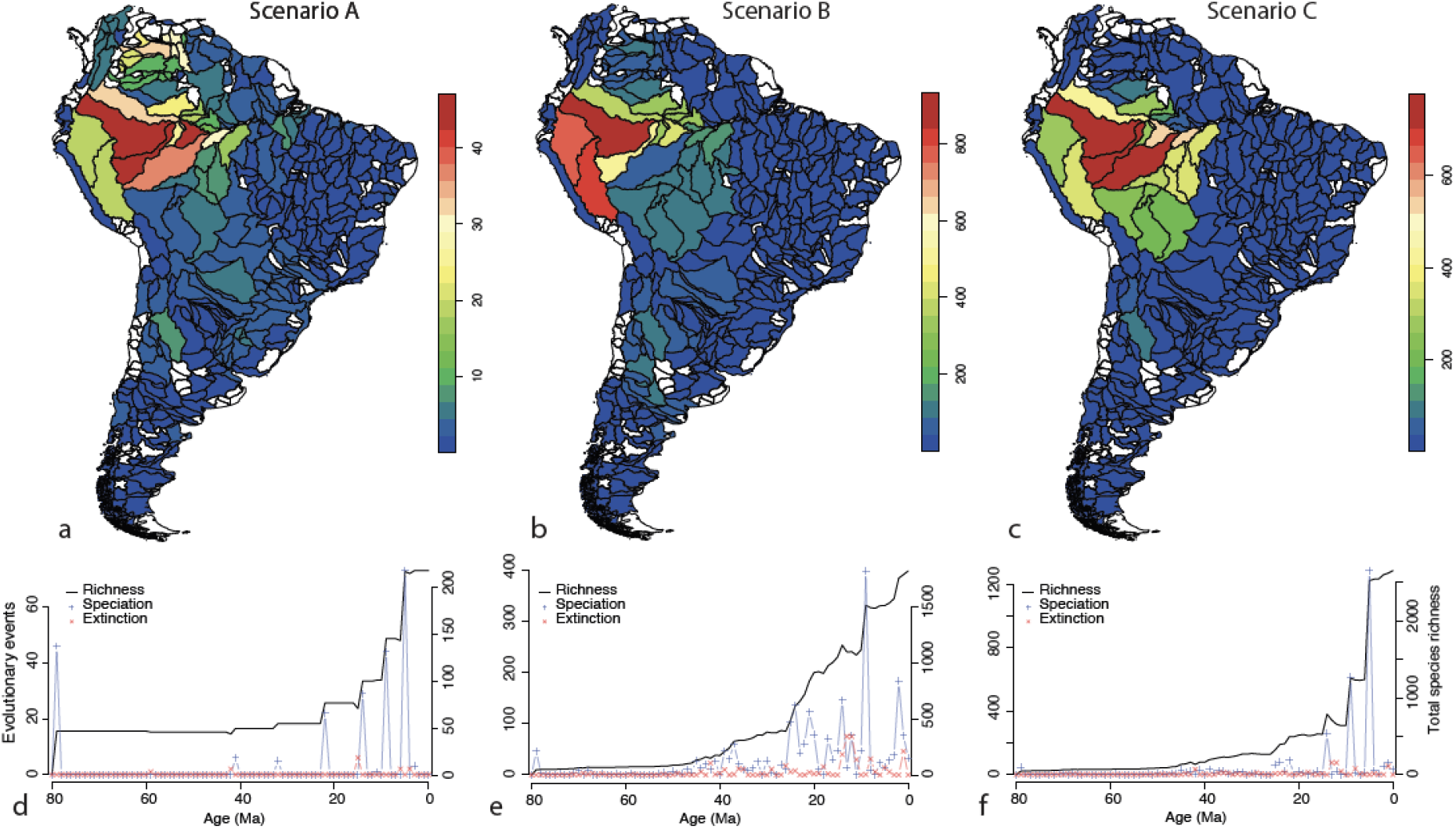
Simulation results. Scenarios as in Fig. 2, *d* = 44.4, *div* = 2, *rs* = 200. **a-c**, Simulated species richness per sub-basin. Comparison with data (Fig. 1b): scenario A: Pearson’s r = 0.54, B: r = 0.57, C: r = 0.61. Basins without large enough rivers (i.e. *rs* < 200 for all cells therein) are depicted in white. **d-f**, Cumulative species richness and speciation and extinction events through model time.

## Geodynamics of landscape evolution

Geodynamic modelling studies have shown that dynamic topography played a role in the formation of the Miocene Pebas wetlands system and the drainage reversal of the Amazon river^23,38,39^. Dynamic subsidence has either been attributed to South America moving westwards over the deep Nazca slab^23^, or to the mid-late Miocene transition from normal subduction to Peruvian flat slab subduction, which was shown to cause subsidence migrating from west to east as the flat slab propagated to the east, consistent with eastward younging ages of the mid-upper Miocene Solimões formation^38^. Based on a combined geodynamic model of surface uplift in the Andes, erosion, and flexure, Sacek^40^ showed that the Miocene drainage reversal may in fact be explained by flexural subsidence alone, but the spatial and temporal extent of the Pebas system can indeed only be explained by a combination of flexural and dynamic subsidence. The conclusion of Sacek^40^, that dynamic topography is not required to explain the drainage reversal of the Amazon, is in line with our findings, which we emphasize are not results of a geodynamic model but instead of a reconstruction based on paleoelevation data, which excludes subsidence in western Amazonia. The age of the drainage reversal produced by the river generation algorithm in this study (10 Ma) coincides well with the age of transcontinental Amazon river formation inferred from sedimentological and paleontological data (9.4 - 9 Ma)^24,25,41,42^.

## Richness gradient along the Amazon river

This study highlights the role of river reorganizations resulting from topographic change in biodiversity dynamics, and shows that frequently-changing river networks result in high speciation rates and promote the accumulation of diversity. So far, patterns of diversity in river networks had primary been explained in a context of present-day (i.e., static) biotic and abiotic conditions^6-8^, yet here we show that the deep-time history of these networks may play a crucial role, particularly in or near tectonically active regions, where river reorganizations are relatively small, but frequent. In contrast to larger river capture events in downstream areas^11^, such minor and frequent river reorganization events will not leave a record in the sedimentary archive detailed enough to reconstruct them, and their thus far underestimated effect can thus only be assessed through reconstruction of river networks and changes therein through a method such as used in this study.

Our modelling results are in line with Oberdorff, et al.^3^, who proposed that the reverse downstream richness gradient in the Amazon river may be explained by historical processes and events, and that the western part of the Amazon Basin has historically been the primary area of species origination. We do however not endorse their conclusion that the richness gradient, which they interpreted to be the result of a west-to-east colonization pattern and an undersaturated eastern Amazonia, can only be explained by a young transcontinental Amazon River that formed within the last 2.5 Ma. First, sedimentological data from exploration wells within the Amazon submarine fan in the Atlantic Ocean indicate that sediments of Andean origin reached the Atlantic coast for the first time between 9.4 and 9 Ma^42^. Second, our modelling results show that even a ∼21 Ma transcontinental Amazon River (scenario B) yields the reverse richness gradient at 0 Ma, and so does a ∼10 Ma transcontinental Amazon River (scenario C). Surface uplift in the northern Andes has been particularly rapid during the Neogene and is still ongoing^22^, with the result that western Amazonia is still the primary area of species origination today. This leads to persistent west-to-east colonization, and a persistent undersaturation of species in eastern Amazonia, despite similar present-day environmental conditions in western and eastern Amazonia.

In this study, we have investigated the effects of spatial changes in fresh water habitat connectivity through geological time, which is just one component of deep-time paleoenvironmental change. Furthermore, we have simplified habitat suitability by assuming that rivers and lakes host the same generic fresh water fish species. Additional work is required to integrate our findings with for example the effects of changes in the provenance area of sediments affecting the geochemistry and nutrient content of rivers, paleoclimate evolution, changes in salinity, and differences in species composition between upland and lowland rivers, and lakes. Nonetheless, our current study is a first step into a mechanistic understanding of diversification and biodiversity evolution through deep geological time.

## METHODS

### Biodiversity data

The data on freshwater fish diversity used in this study is derived from Cassemiro, et al.^4^. These authors collected occurrence data from 4967 freshwater fish species in South America, and calculated species richness values for all drainage sub-basins (490 basins; level 5 basins of the HydroBASINS database^17^). The occurrence dataset revealed a disbalance in sampling effort between the different basins, with basins of comparable size containing occurrence records varying by a factor ten. To account for this sampling heterogeneity across the basins, Cassemiro, et al.^4^ performed a correction procedure consisting of (1) calculating a ‘completeness index’^43^, indicating the probability that adding an additional occurrence record would add a new species to a basin; (2) in case of a completeness index lower than 0.75, randomly sampling occurrence records from adjacent basins to increase the number of occurrences and species; and (3) iteratively repeating this process, until all basins have a completeness index higher than 0.75. This procedure was repeated 100 times, after which the average richness per basin was calculated. Twenty poorly sampled basins were excluded.

### Paleogeographic reconstruction

The dynamic landscape used as input in the biodiversity simulations is based on the reconstruction of Andean mountain building of Boschman^22^. Boschman^22^ compiled estimates of paleoelevation and surface uplift for 36 individual domains in the Andes, and developed a reconstruction of 80 million years of paleoelevation in 1 Ma, 0.1° resolution. This reconstruction is for the Andes only and does not include estimates of paleoelevation of the rest of South America. We assume that the current topography in the ancient Guianan and Brazilian shield areas was already present at 80 Ma. However, we consider the relatively high elevations present along the eastern slopes of the Andes to be the result of the deposition of erosional material derived from the Andes during mountain building, and we reconstruct them as such, by lowering their elevation synchronously with the Andean mountain ranges to the west (Fig. S6), using the topography reconstruction method of Boschman^22^. We include the Miocene Pebas system and smaller preceding and following lakes, wetlands, and marine incursions, based on the maps of Hoorn and Wesselingh^20^ for northern South America and Hernández, et al.^44^ for southern South America (Fig. S7).

### River generation from paleoDEMs

From the paleoDEMs, we generated drainage networks via a method derived from the D8 algorithm^45^ and based on code developed for the *OCNet* R-package^46^. First, drainage directions are established based on the steepest descent between neighboring grid cells. A cell is not attributed a drainage direction if all neighboring cells have equal (i.e. the cell belongs to a flat area) or higher elevations (i.e. the cell is an internal outlet). Second, drainage directions for flat areas are attributed by following the algorithm of Barnes, et al.^47^, which produces drainage directions away from higher terrain and towards lower terrain. Third, to solve for internal outlets, we apply an iterative procedure: for each internal outlet *o* (sorted by decreasing elevation), the contour of the region draining into *o* is determined, and the grid cell *c* on that contour with the lowest elevation is determined. The drainage path from *c* towards *o* is then reversed, so that *c* becomes the outlet of the catchment. If *c* borders a cell *c’* that drains towards an outlet (either internal or along the coastline of the continent) whose elevation is lower than that of *o*, then a drainage direction from *c* to *c’* is established; if more than one such cell *c’* exist, the one draining towards the outlet with lowest elevation is selected. This procedure is repeated until all outlets coincide with the shoreline of the South American continent. To avoid the algorithm ‘finding’ the modern incised river valleys for each of the 80 time steps, we remove these present day features from the landscape by lifting up all land lower than 100 m meter above sea level to 100 m. This approach proves robust as the algorithm produces river networks for the present day that match the modern configuration of South American rivers well (Fig. 2, 4). The obtained drainage networks include drainage directions for each individual 0.1° grid cell in South America, as well as a value for drainage area (i.e. how many cells drain into the cell). The resolution of the river network used in the gen3sis simulations is controlled through this parameter *river size* (*rs*), thus using drainage area as a proxy for stream size^9^.

**Fig. 4.**
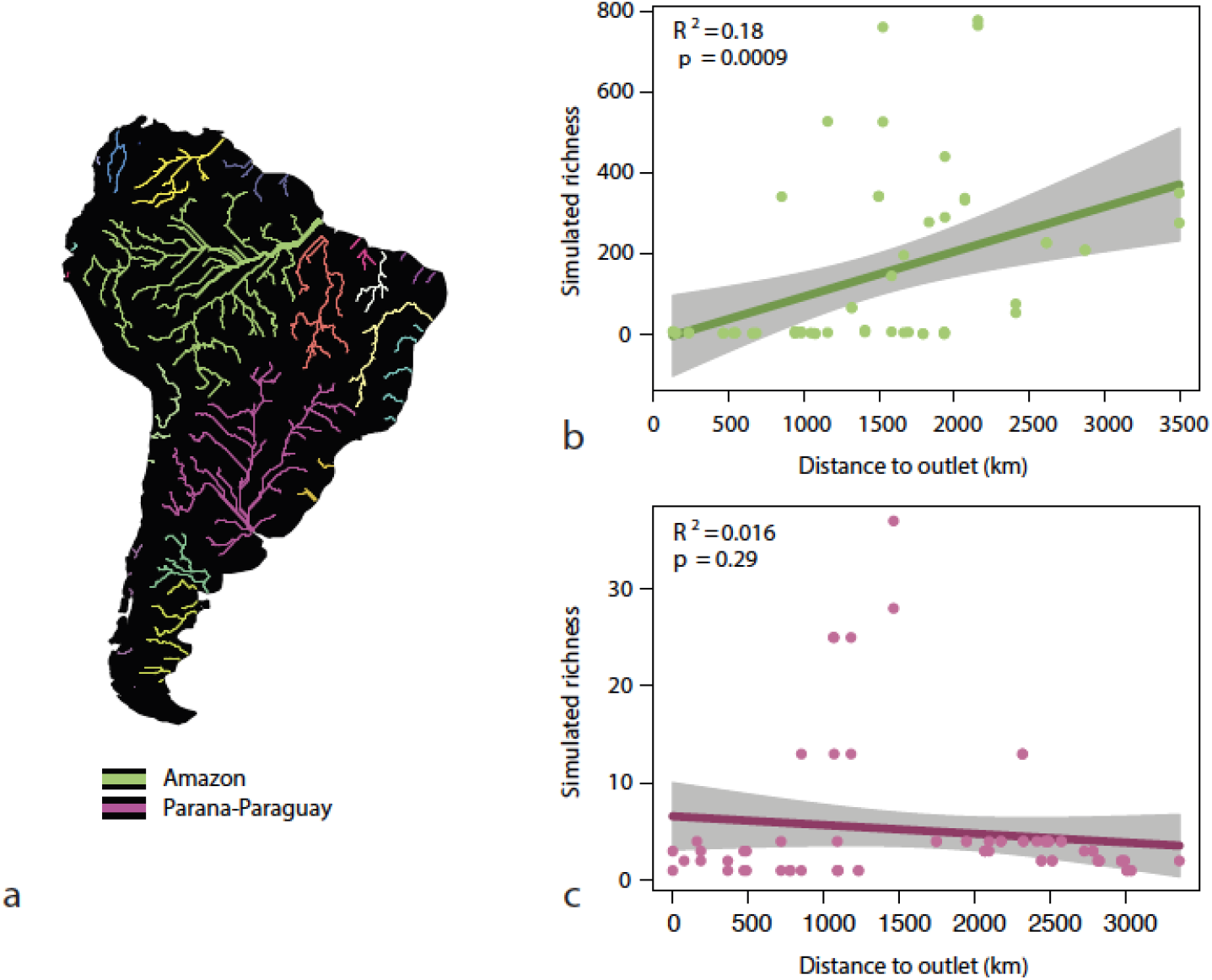
a, River reconstruction algorithm results. b and c, Simulated richness versus distance to outlet for the sub-basins in the Amazon and Parana-Paraguay river basins. The positive relationship for the Amazon river shown in Fig. 1c is reproduced by the model, suggesting that the paleogeographic history of South America is the primary driver of the richness gradient along the Amazon river. For comparison, there is no significant correlation between simulated richness and distance to outlet for the Parana-Paraguay river, illustrating the absence of abiotic and biotic environmental factors in the modelling approach, which are expected to shape richness patterns with a negative relationship between richness versus distance to outlet (see Fig. 1d).

### Gen3sis modelling

Gen3sis (GENeral Engine for Eco-Evolutionary SImulationS^16^) is a mechanistic model simulating the evolution of biodiversity as a function of habitat change, through the mechanisms of dispersal, speciation, and extinction. Habitat change is characterized by environmental conditions related to landscape evolution in a geographical framework, and is in this study represented by changes in suitable (rivers, lakes) or non-suitable (land, marine) habitat. The simulations are initiated with a single species that is present in all suitable habitat cells at 80 Ma, in line with the Late Cretaceous origin of the extant South American fish fauna^10,12,13^. Each (one million year) time step, species disperse to all connected suitable habitat cells determined by a dispersal rate *d*. For each time step that two populations of the same species are disconnected, the populations build up ‘genetic distance’, and once this genetic distance reaches the divergence threshold (*div*), the two populations become two different species. A species goes extinct when all cells in its range change from suitable to non-suitable habitat. The model used in this study uses three input parameters: the minimum river size (*rs*), the divergence threshold (*div*), and the dispersal rate (*d*). The river size and divergence threshold parameters control the spatial and temporal resolution of speciation, which primarily affects the number of species generated by the model, but not species richness patterns (Fig. S2, S3). Therefore, we set these parameters to the finest resolution that allowed for a viable computation time (*rs* = 200, *div = 2*). The dispersal rate affected both the number of species generated by the model and the richness pattern of these species. We therefore simulated each scenario for a range of dispersal rates (*d* = 22.2, 33.3, 44.4, 55.5, 66.6, 77.7, and 88.8 km/Myr, which roughly correspond to 2, 3, 4, 5, 6, 7, and 8 grid cells/Myr, Fig. S4) and compared values of simulated species richness to observed species richness per catchment by calculating a Pearson’s r, thereby assessing richness patterns and not absolute species numbers. In Fig. 3, we report the simulations in which the result of scenario C (the scenario that most accurately portrays the paleographic history of South America) matches the data best (*d* = 5 (i.e., 55.5 km/Myr); r = 0.61, Fig. S4).

## Acknowledgements

This work was funded by ETH postdoctoral fellowship 18-2 FEL-52 granted to LMB, and Swiss National Science Foundation Grants PP00P3_179089 granted to FA and 31003A_173074 granted to LP.

## Author Contributions

LMB: Analysis, visualization, writing; FASC, LC, JdV: methodology; review and editing, FA, LP: conceptualization, review and editing; OH, CH: review and editing.

