## Supplementary Materials for "South American freshwater fish diversity shaped by Andean uplift since the Late Cretaceous"

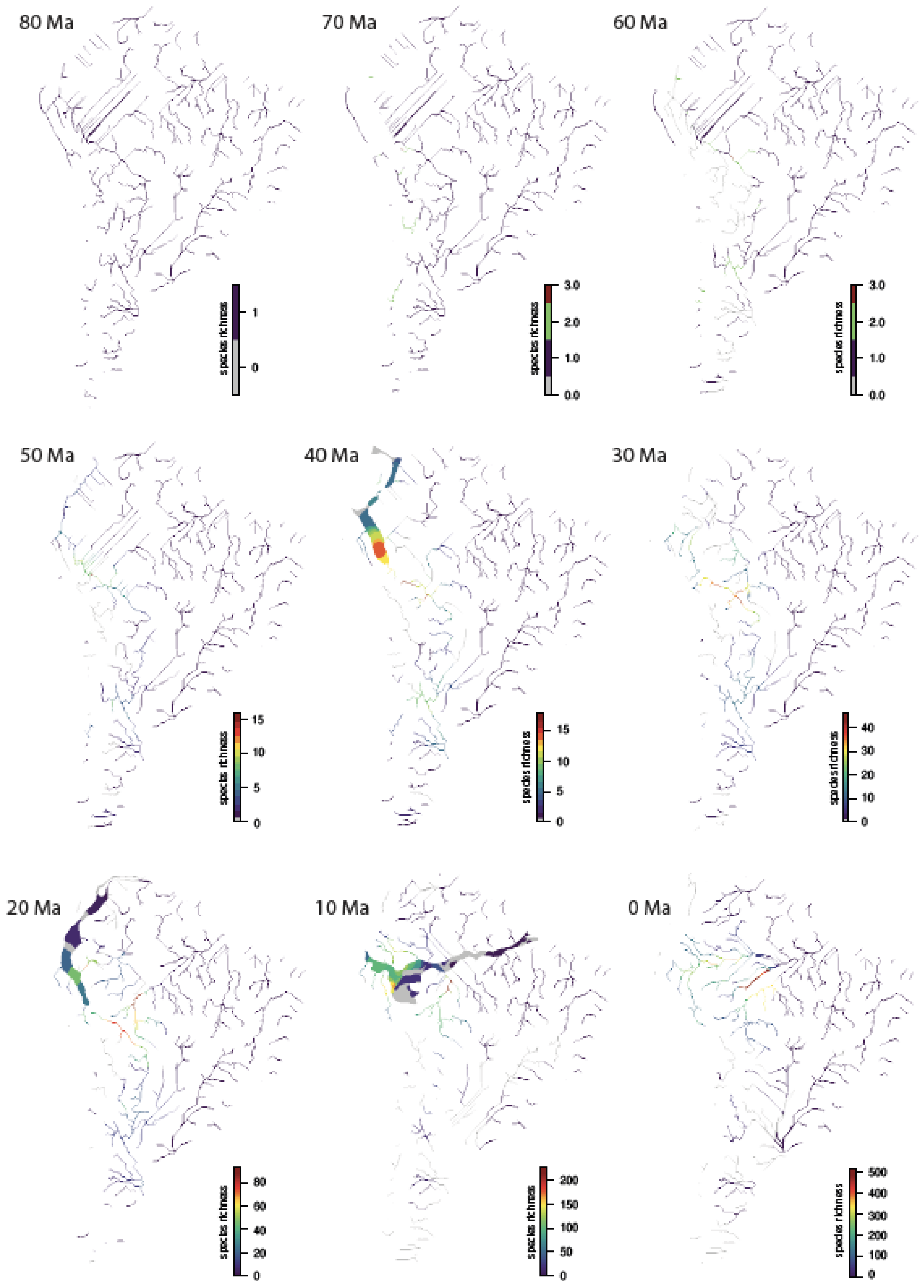


**Fig. S1 | Example of a gen3sis simulation.** Snapshots at 80, 70, 60, 50, 40, 30, 20, 10 and 0 Ma; results of richness per sub-basin shown in Fig. 3c.

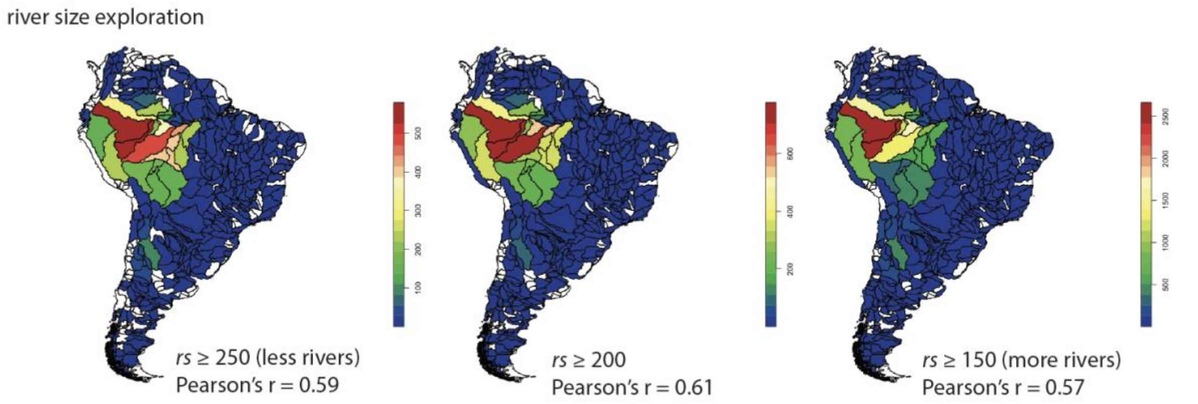


**Fig. S2 | Parameter exploration: river size (*rs*).** *d* = 5, *div* = 2.

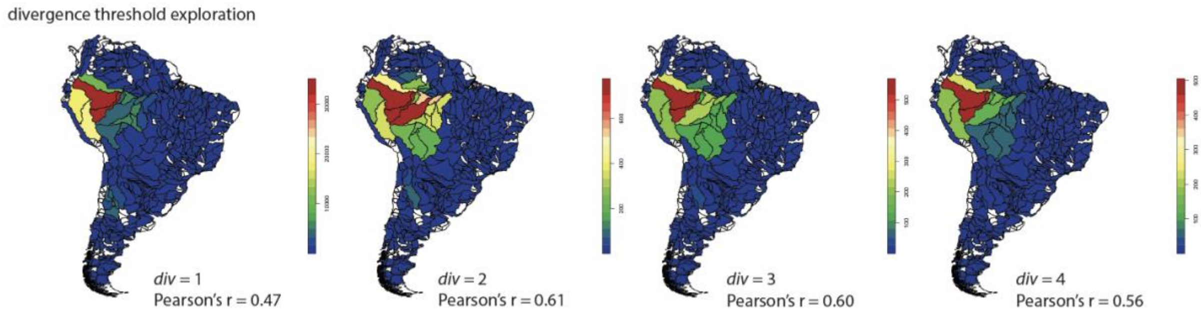


**Fig. S3 | Parameter exploration: divergence threshold (*div*).** *d* = 5, *rs* = 200.

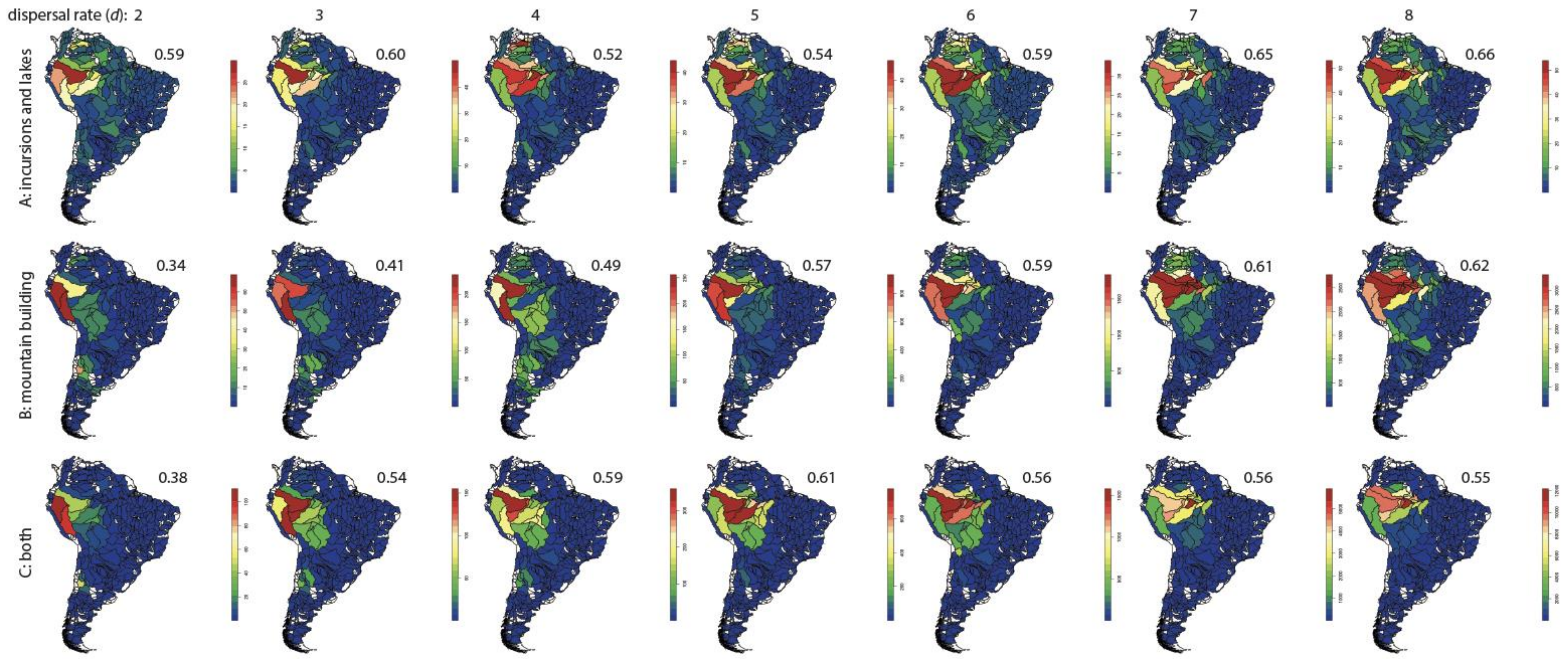


**Fig. S4 | Parameter exploration: dispersal rate (*d*).** *div* = 2, *rs* = 200. Numbers are Pearson's r.

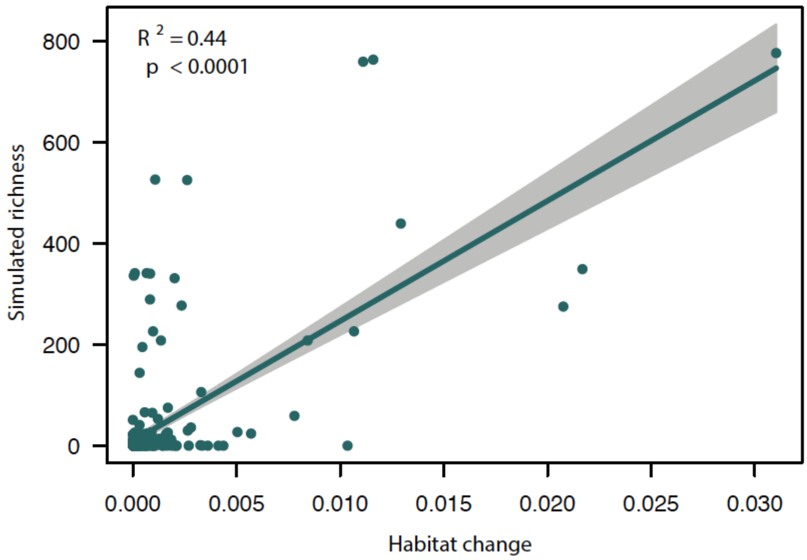


**Fig. S5 | Simulated richness versus cumulative habitat change of sub-basins.** Cumulative habitat change per sub-basin (shown in Fig. 2) is a measure of the amount of cells changing from suitable to unsuitable habitat, or vice versa, summed for all 80 time steps.

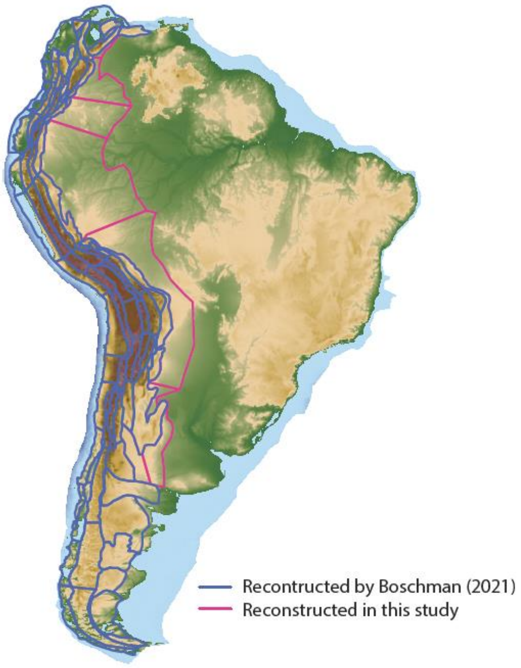


**Fig. S6 | Map of South America, outlining the area in which elevation is reconstructed by the study of Boschman[1], and this study.**

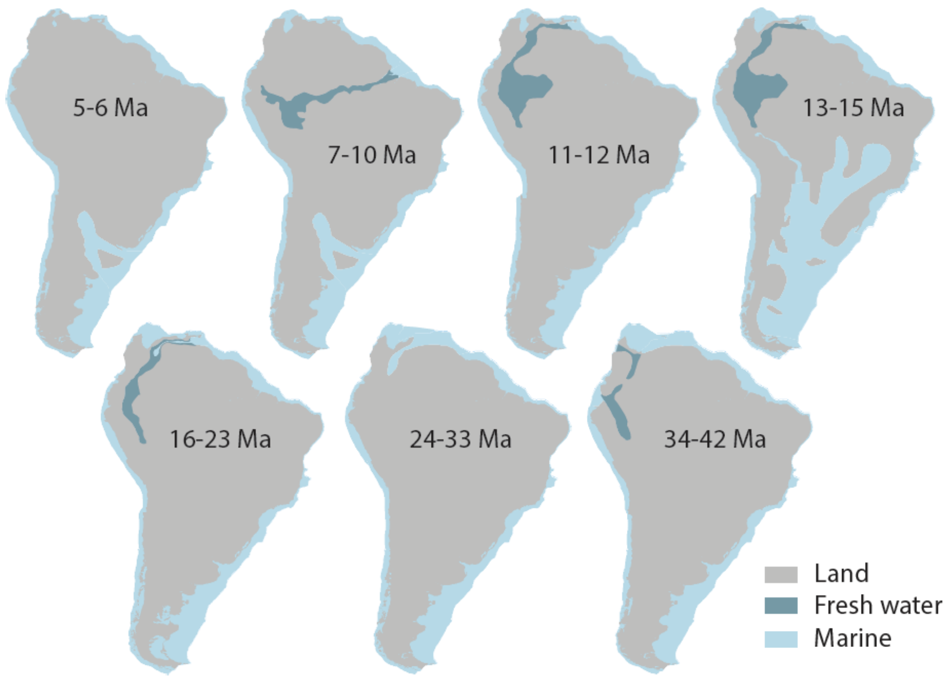


**Fig. S7 | Outlines of lakes and marine incursions included in the reconstruction, based on the maps of Hoorn and Wesselingh[2] for northern South America and Hernández, et al.[3] for southern South America.**

1 Boschman, L. M. Andean mountain building since the Late Cretaceous: A paleoelevation reconstruction. *Earth-Science Reviews*, doi:10.1016/j.earscirev.2021.103640 (2021).

2 Hoorn, C. & Wesselingh, F. *Amazonia: landscape and species evolution: a look into the past*. (John Wiley & Sons, 2011).

3 Hernández, R. *et al.* Age, distribution, tectonics, and eustatic controls of the Paranense and Caribbean marine transgressions in southern Bolivia and Argentina. *Journal of South American Earth Sciences* **19**, 495-512 (2005).
